# Identification and validation of aging-related genes in heart failure based on multiple machine learning algorithms

**DOI:** 10.1101/2023.12.05.570061

**Authors:** Yiding Yu, Lin Wang, Wangjun Hou, Yitao Xue, Xiujuan Liu, Yan Li

## Abstract

**Background:** In the face of continued growth in the elderly population, the need to understand and combat age-related cardiac decline becomes even more urgent, requiring us to uncover new pathological and cardioprotective pathways.

**Methods:** We obtained the aging-related genes of heart failure through WGCNA and CellAge database. We elucidated the biological functions and signaling pathways involved in heart failure and aging through GO and KEGG enrichment analysis. We used three machine learning algorithms: LASSO, RF and SVM-RFE to further screen the aging-related genes of heart failure, and fitted and verified them through a variety of machine learning algorithms. Finally, we searched for drugs to treat age-related heart failure through the DSigDB database.

**Results:** We obtained 57 up-regulated and 195 down-regulated aging-related genes in heart failure through WGCNA and CellAge databases. GO and KEGG enrichment analysis showed that aging-related genes are mainly involved in mechanisms such as Cellular senescence and Cell cycle. We further screened aging-related genes through machine learning and obtained 14 key genes. We verified the results on the test set and 2 external validation sets using 15 machine learning algorithm models and 207 combinations, and the highest accuracy was 0.911. Through screening of the DSigDB database, we believe that rimonabant and lovastatin have the potential to delay aging and protect the heart.

**Conclusions:** We identified aging signature genes and potential therapeutic drugs for heart failure through bioinformatics and multiple machine learning algorithms, providing new ideas for studying the mechanism and treatment of age-related cardiac decline.

## INTRODUCTION

With an aging global population and improved survival rates for ischemic heart disease due to increasingly effective, evidence-based treatments, heart failure prevalence is on the rise, now affecting over 64 million individuals worldwide^1^. This trend is particularly pronounced among elderly patients. As such, the escalating elderly demographic intensifies the urgency to both comprehend and counteract age-related cardiac deterioration. This necessitates the exploration of novel pathological and cardioprotective mechanisms, aiming to reduce the extensive impact on global public health^2^.

Heart failure’s development is intricately linked to the complex interplay of cardiovascular aging, risk factors, comorbidities, and disease moderators^3^. While dietary restrictions, increased physical activity, and pharmacological interventions are pivotal in decelerating cardiovascular function decline in aging populations, their impact on mortality remains limited^4,5^. Recent research posits that the persistent high prevalence of cardiovascular diseases and associated mortality may stem from a lack of targeted interventions addressing the aging process directly. These studies have identified eight key molecular markers characteristic of cardiovascular aging: impaired macroautophagy, proteostasis loss, genomic instability, epigenetic changes, mitochondrial dysfunction, cellular senescence, disrupted neurohormonal signaling, and inflammation^6^.

With advancing age, significant structural and functional transformations occur in the heart, blood vessels, and microcirculation^7^. These changes in cardiac structure and function contribute to an increased vulnerability to heart failure in the elderly. However, the precise mechanisms by which aging precipitates heart failure are not yet fully understood. Unraveling the specific genes and molecular processes involved in the onset and progression of heart failure during aging is crucial. Such insights are expected to pave the way for innovative strategies to combat age-related decline, preserve circulatory function, and extend the disease-free lifespan of individuals.

Bioinformatics, an ever-evolving multidisciplinary domain, is revolutionizing our understanding in the medical sciences. This study leverages high-throughput technologies and machine learning to unearth pivotal aging genes and molecular pathways implicated in heart failure. Utilizing Weighted Gene Co-expression Network Analysis (WGCNA), we identified crucial module genes from the largest available heart failure dataset, integrating these findings with the CellAge database to highlight aging-related genes. Subsequent feature enrichment analysis led to the selection of three advanced machine learning algorithms: Least Absolute Shrinkage and Selection Operator (LASSO), Random Forest (RF), and Support Vector Machine Recursive Feature Elimination (SVM-RFE), for pinpointing key genes associated with heart failure and aging. To ascertain the robustness of our findings, we employed a comprehensive validation approach, testing the results across a primary test set and two external datasets using 15 distinct machine learning models and 207 unique combinations. In the final phase of our study, we conducted a comprehensive drug prediction analysis utilizing the Drug Signatures Database (DSigDB). This approach was instrumental in identifying potential pharmaceutical candidates for the management and treatment of heart failure and associated aging processes. The methodology and progression of this study are encapsulated in Figure 1, which outlines the research flowchart.

**Figure 1:**
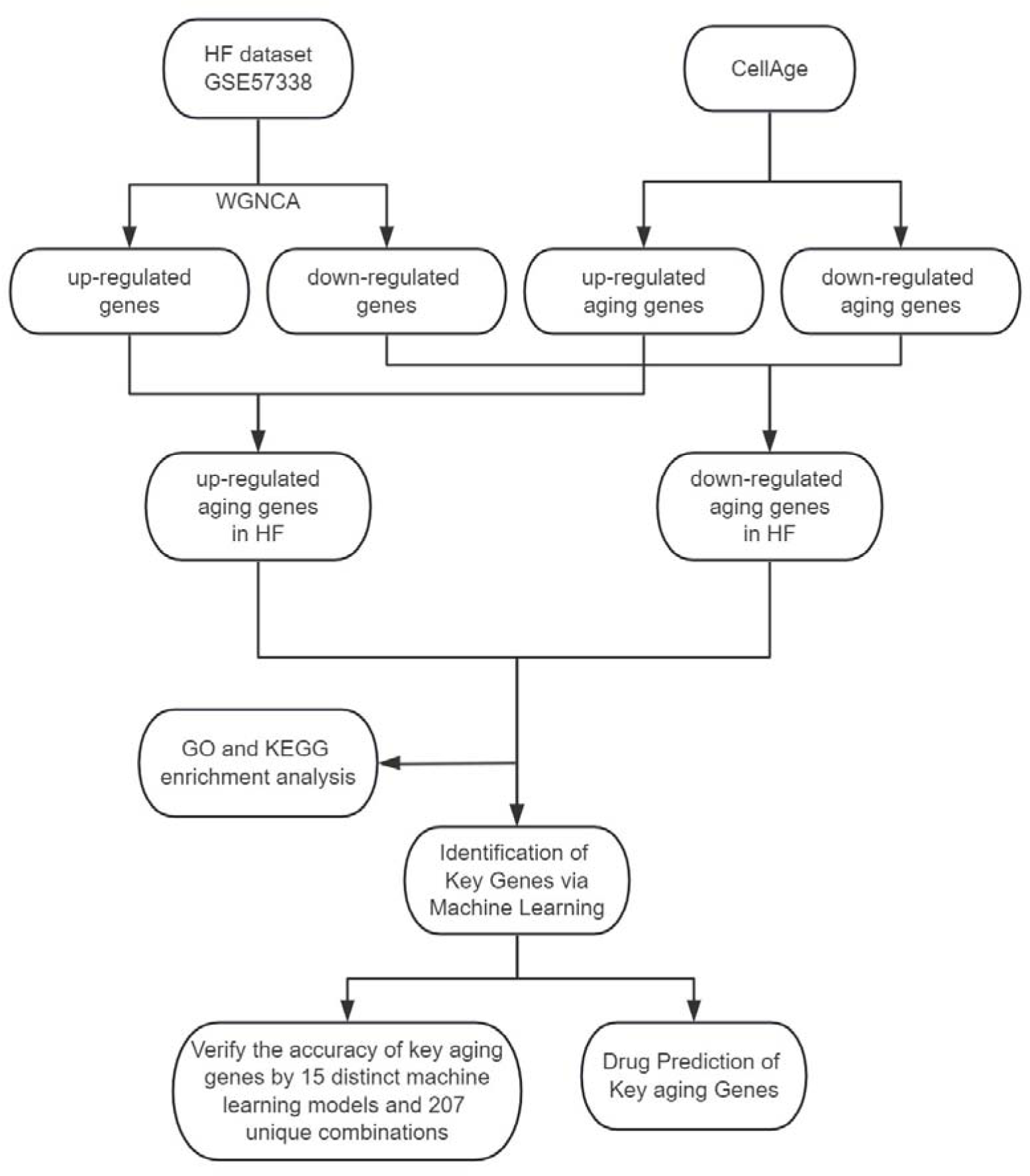
The study flowchart.

## MATERIALS AND METHODS

### Data Acquisition and Preprocessing

This study’s data were sourced from the publicly accessible Gene Expression Omnibus (GEO) database, with the datasets having previously obtained participant consent and ethical approval^8^. Consequently, our research did not require additional approval from an institutional review board. We selected GSE57338 as our primary dataset for ischemic heart failure analysis due to its extensive sample size, comprising left ventricular myocardial samples from 95 ischemic heart failure patients and 136 individuals without heart failure^9^. For external validation, we utilized datasets GSE5406 (including myocardial samples from 108 ischemic heart failure patients and 16 non-heart failure individuals) and GSE16499 (comprising samples from 15 ischemic heart failure patients and 15 non-heart failure individuals)^10,11^.

Data preprocessing was conducted using R software (version 4.2.0). In this process, we eliminated probes linked to multiple molecules. Where multiple probes corresponded to a single molecule, only the probe with the highest signal value was retained. To ensure data consistency and accuracy, we also corrected for batch effects in the data and converted probe IDs to gene symbols based on the platform’s annotation file.

### Weighted Gene Co-expression Network Analysis

We used the WGCNA package to explore gene modules associated with heart failure^12^. Using 0.5 as the filtering standard and removing unqualified genes and samples through the goodSamplesGenes function, a scale-free co-expression network was established. Subsequently, adjacency was calculated with a default soft threshold of β = 30 and scale-free R2 = 0.9, and the adjacency was converted into a topological overlap matrix (TOM) to determine gene ratios and dissimilarity. Genes with the same expression profile are divided into gene modules using average linkage hierarchical clustering, we prefer larger modules, so we set the minimum module size to 300. Finally, the dissimilarity of module characteristic genes is calculated, the cutting line of the module dendrogram is selected to combine several modules for further research, and the visualization of the characteristic gene network is completed.

### Screening Candidate Aging-related Genes in HF

The CellAge database (https://genomics.senescence.info/cells/) serves as a comprehensive repository of human genes associated with cellular senescence^13^. This database meticulously catalogs genes with established positive, negative, or undetermined impacts on this process.We involved correlating genes upregulated in heart failure (HF) with those in CellAge known to accelerate cell senescence. Concurrently, we analyzed the overlap of genes downregulated in HF with those identified in CellAge as senescence inhibitors. This dual-faceted approach facilitated the identification of key aging-related genes specifically involved in the pathophysiology of heart failure.

### Functional Enrichment Analysis

To elucidate the biological processes and functions of aging genes implicated in heart failure, our study utilized the clusterProfiler package^14^. This tool enabled us to conduct a comprehensive Gene Ontology (GO) and Kyoto Encyclopedia of Genes and Genomes (KEGG) enrichment analysis^15,16^. Through this analysis, we were able to identify and visualize key pathways and gene functions, providing deeper insights into how aging genes contribute to the pathophysiology of heart failure.

### Machine Learning

In our investigation, we employed three distinct machine learning algorithms—LASSO, RF, and SVM-RFE—to rigorously identify key aging genes in HF^17–19^. The LASSO algorithm was executed using the glmnet package, incorporating ten-fold cross-validation to pinpoint significant genes. For the RF algorithm, we utilized the randomForest package, selecting the top 20 genes as our primary candidates. The SVM-RFE algorithm, conducted via the e1071 package, was used to determine the optimal gene subset based on accuracy. The culmination of these methodologies was the identification of a consensus set of genes, representing the intersection of results from all three algorithms, which we designated as the critical aging genes in heart failure.

### Validation of Key Aging Genes

In order to verify the accuracy of key aging genes, we integrated 15 machine learning algorithms (included Neural Networks, Logistic Regression, Linear Discriminant Analysis, Quadratic Discriminant Analysis, K-Nearest Neighbors, Decision Trees, Random Forest, XGBoost, Ridge Regression, LASSO Regression, Elastic Net Regression, Support Vector Machines, Gradient Boosting Machines, Stepwise Logistic Regression, and Naive Bayes) and combined these 15 algorithms through caret parameter adjustment, custom parameter combination, lasso feature screening, and cross-validation, resulting in a total of 207 machines learning model. For our analysis, we randomly allocated 70% of the GSE57338 dataset as the training set and designated the remaining 30% for testing. In addition, we incorporated two external validation sets, GSE5406 and GSE16499, to further strengthen the robustness of our results.

### Drug Prediction

DSigDB, a comprehensive drug signature database, was employed for gene set analysis in our study^20^. We utilized the identified key aging genes as a reference list, applying DSigDB’s predictive capabilities to identify potential drug molecules.

## RESULTS

### Construction of Co-expressed Gene Modules

In our study, we conducted WGCNA on the GSE57338 dataset, identifying both upregulated and downregulated modules significantly associated with HF. The analysis revealed that a β value of 10 brought the network closest to a scale-free topology. Within this framework, we pinpointed 9 modules related to HF. Notably, in the upregulated category, the pink module (correlation coefficient = 0.68, P = 2e-32) and the green module (correlation coefficient = 0.56, P = 1e-20) demonstrated the highest correlation with HF, encompassing a combined total of 2,998 genes. Conversely, among the downregulated modules, the turquoise module exhibited the strongest association with HF (correlation coefficient = 0.48, p = 2e-14), comprising 7,570 genes. These findings are represented in Figure 2-A and Figure 2-B.

**Figure 2:**
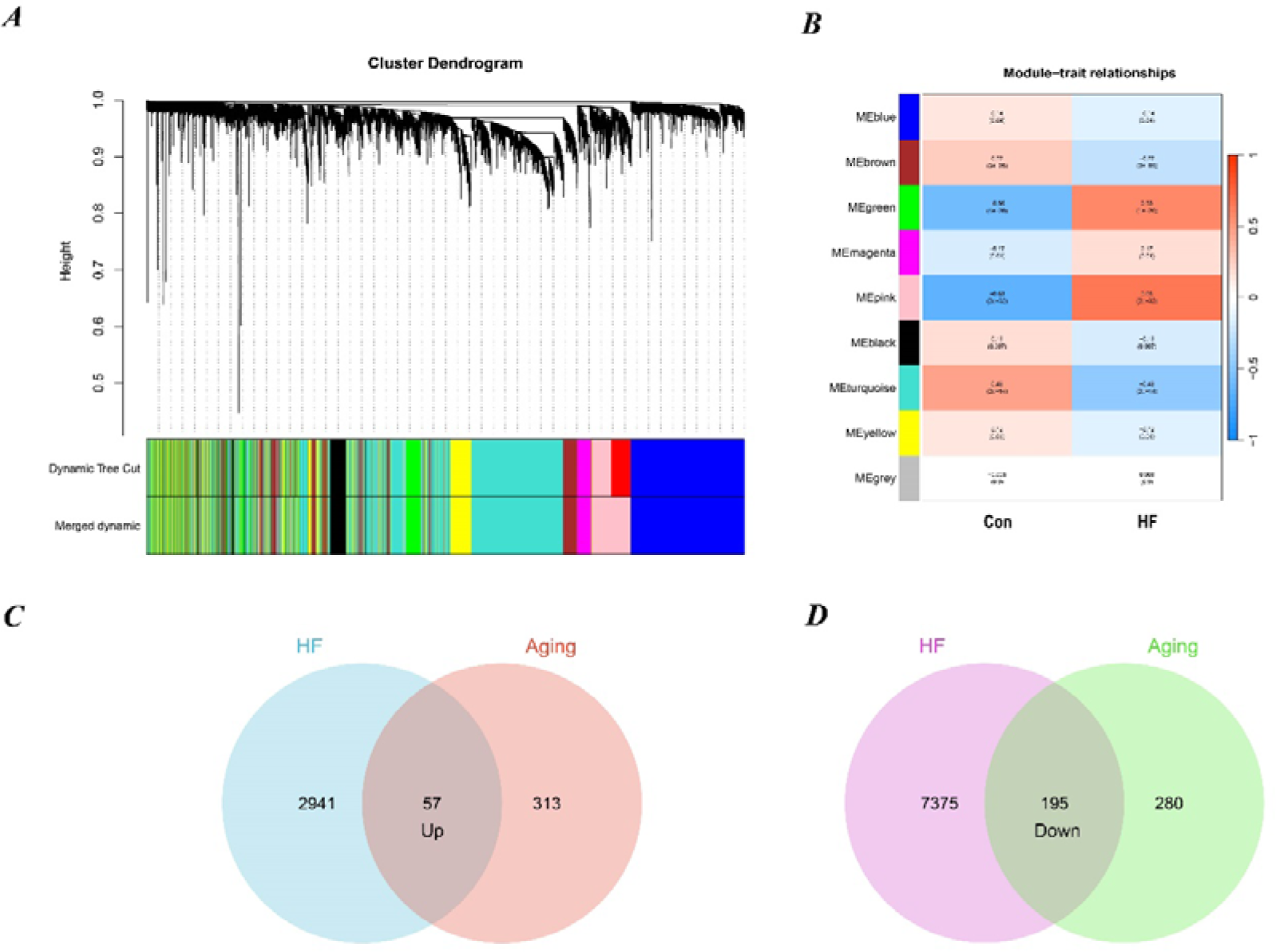
Identification of aging-related genes. (A) Gene and trait clustering dendrograms of HF. Gene clustering trees (dendrograms) obtained by hierarchical clustering of neighbor-based differences. (B) 9 gene co-expression modules of HF. The numbers in each cell means the correlation coefficient and p-value. (C) 57 genes promote both aging and HF. (D) 195 genes inhibit both aging and HF.

From the CellAge database, we identified 370 genes implicated in promoting aging and 475 genes associated with inhibiting aging. We then conducted an intersection analysis between these aging-related genes and those influencing heart failure. This approach revealed 57 genes that concurrently promote both aging and heart failure. We also identified 195 genes that play a role in inhibiting both aging and heart failure. The results of these intersection analyses are represented through Venn diagrams in Figure 2-C and Figure 2-D.

### Functional Enrichment Analysis

We performed GO and KEGG enrichment analysis on 252 aging-related genes in heart failure. This was undertaken to elucidate the shared biological mechanisms underpinning both conditions. The GO enrichment analysis encompassed three primary categories: Biological Process (BP), Cellular Component (CC), and Molecular Function (MF). Notably, BP categories were predominantly focused on aspects like histone modification, regulation of the mitotic cell cycle, cell cycle phase transition, and positive regulation of cell division. CC categories emphasized elements such as focal adhesion, cell-substrate junctions, pericentric heterochromatin, SWI/SNF superfamily-type complexes, and ATPase complexes. In the MF category, significant functions included histone binding, DNA-binding transcription factor interaction, transcription coregulator and corepressor activities, and NAD-dependent histone deacetylase activity.

The KEGG enrichment analysis revealed that these aging-related genes in heart failure were significantly enriched in pathways including Cellular Senescence, Proteoglycans in Cancer, Cell Cycle, MicroRNAs in Cancer, C-type Lectin Receptor Signaling Pathway, PI3K-Akt Signaling Pathway, and various cancer-related signaling pathways. The top five results from the GO enrichment analysis and the top ten from the KEGG enrichment analysis will be presented, as depicted in Figure 3.

**Figure 3:**
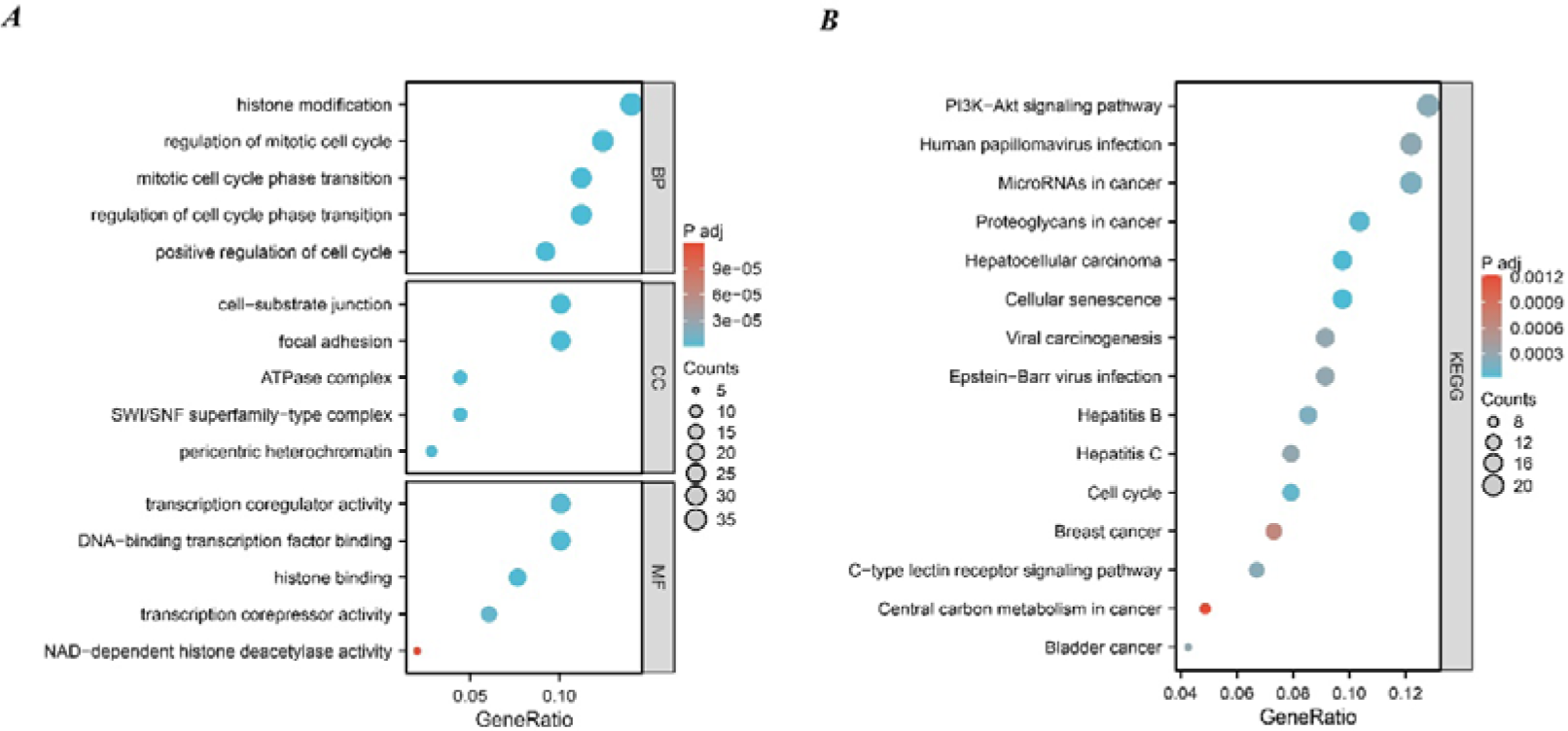
Function enrichment analysis of 252 aging-related genes. (A) GO enrichment analysis results. (B) KEGG enrichment analysis results.

### Identification of Key Genes via Machine Learning

We used three machine learning algorithms, LASSO, RF and SVM-RFE, to further screen key aging genes in heart failure. Among the up-regulated genes, the LASSO algorithm identified 17 candidate genes. The RF algorithm ranks genes according to the importance calculation of each gene, and we select the top 20 genes as candidate genes. The SVM-RFE algorithm shows that the accuracy is highest when 34 genes are included, so we selected the first 34 genes of the SVM-RFE algorithm as candidate genes. After intersecting the results of the three algorithms, we obtained 10 up-regulated key aging genes in heart failure, namely CDKN1B, SPIN1, GNMT, HTRA1, ITPK1, MAVS, MME, RAF1, TLR3, and XAF1.

Similarly, among the down-regulated genes, the LASSO algorithm identified 39 candidate genes. We still select the top 20 genes of the RF algorithm as candidate genes. The SVM-RFE algorithm shows that the accuracy is highest when 14 genes are included, so we selected the first 14 genes of the SVM-RFE algorithm as candidate genes. After intersecting the results of the three algorithms, we obtained four down-regulated key aging genes in heart failure, namely BCL6, EIF4EBP1, MEIS2, and SMARCA2. The visualization results are shown in Figure 4.

**Figure 4:**
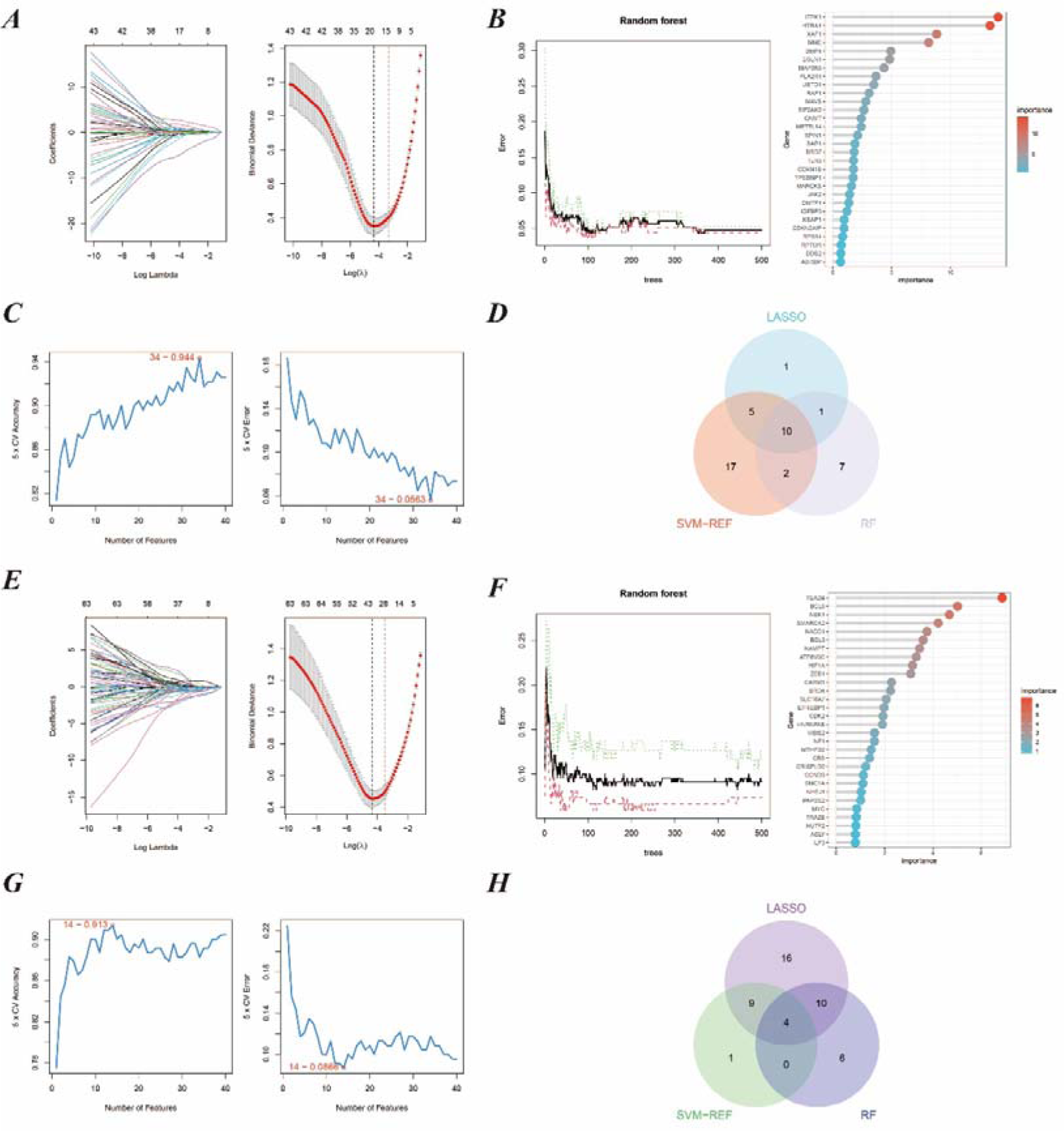
Machine learning in screening key aging genes for HF. (A) Screening of key aging genes using the Lasso Model in up-regulated genes. The Lasso coefficient profiles were utilized to identify the optimal feature genes, with the optimal lambda determined by minimizing the partial likelihood deviance. Each coefficient curve in the left picture represents an individual gene. The solid vertical lines in the right picture represent the partial likelihood deviance, and the number of genes (n = 17) corresponding to the lowest point of the curve was deemed most suitable for the Lasso model. (B) Screening of key aging genes using the RF Model in up-regulated genes. The relative importance of overlapping candidate genes was calculated using the random forest approach. We present the results for the top 20 genes. (C) Screening of key aging genes using the SVM-RFE Model in up-regulated genes. The SVM-RFE algorithm was employed to further identify the optimal feature genes, based on the highest accuracy and lowest error obtained from the curves. The x-axis indicates the number of feature selections, while the y-axis represents the prediction accuracy. (D) Venn diagram illustrating the identification of 10 candidate genes for up-regulated genes through the aforementioned three algorithms. (E) Screening of key aging genes using the Lasso Model in down-regulated genes. (F) Screening of key aging genes using the RF Model in down-regulated genes. (G) Screening of key aging genes using the SVM-RFE Model in down-regulated genes. (H) Venn diagram shows that 4 key aging genes for down-regulated genes are identified via the above three algorithms.

### Key Genes Verification

We identified 14 key aging genes associated with heart failure using three distinct machine learning algorithms. To circumvent the limitations imposed by the sample size in the ROC curve analysis of combined genes, we pursued an alternative validation strategy. Specifically, we validated these 14 genes using 15 different machine learning algorithm models and 207 combinations, across both the test set and two external validation sets. The validation results indicated that, in most algorithms, the accuracy of these 14 genes exceeded 0.8 in both the test set and external validation sets. Notably, the Elastic Net Regularized Generalized Linear Model with Cross-Validation (ENR-CV), with specific parameters set to 10-fold cross-validation, a cutoff value of 0.25, and an alpha value of 0.6, achieved the highest average accuracy (0.911). The top 50 average accuracy rankings from this comprehensive analysis are depicted in Figure 5.

**Figure 5:**
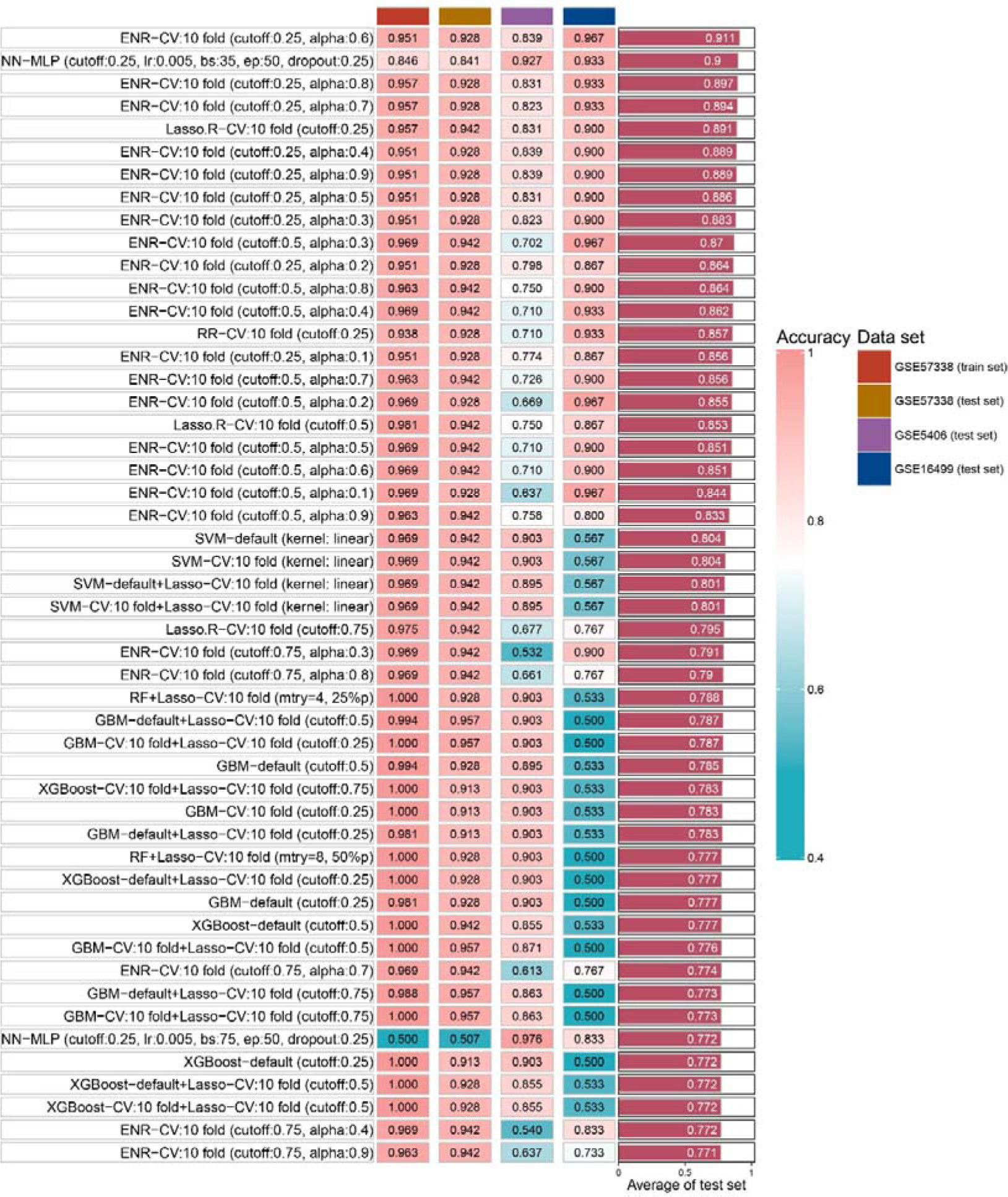
The accuracy of top 50 machine-learning algorithm combinations.

### Drug Prediction of Key Genes

In our pursuit to identify potential pharmacological agents for tackling heart failure in aging patients, we utilized the DSigDB database. Our selection criteria focused on drugs with an Adjusted P-value of less than 0.01. This stringent threshold led to the identification of six promising drug candidates: Arsenenous acid, Cyclophosphamide, Lovastatin, Rimonabant Hydrochloride, Sorafenib, and Alvespimycin. Detailed information about these drugs was provided in Table 1.

**Table 1:**
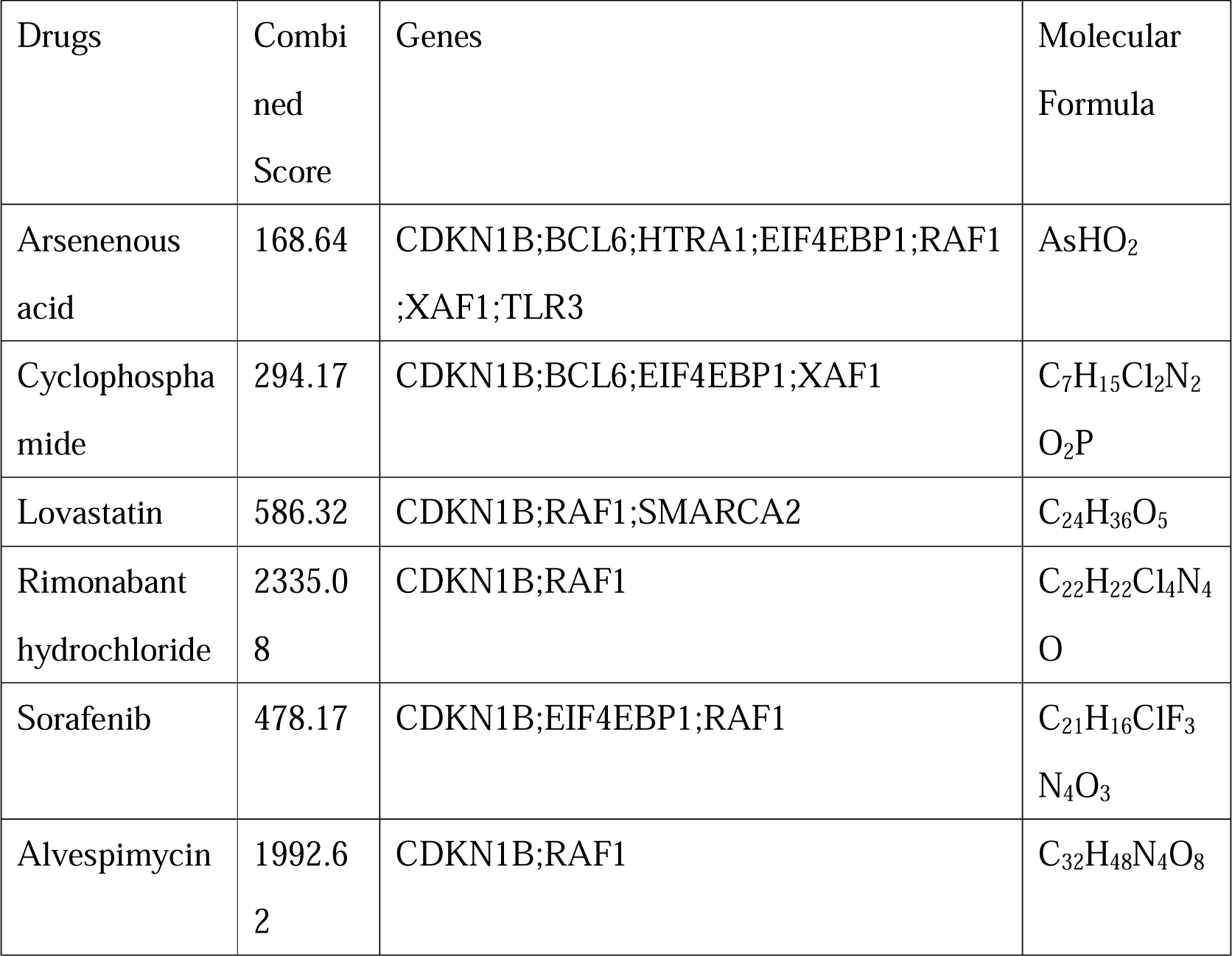
The details of drugs.

## DISCUSSION

Heart failure is a disease caused by structural changes or functional impairment of the heart, and aging plays an important role in its progression^21^. In fact, among the elderly, maintaining normal circulatory function helps increase disease-free life expectancy and maintain a higher quality of life^22^. As the aging of the population increases and the survival rate of ischemic cardiomyopathy increases, heart failure in the elderly population has brought serious economic and public health burdens^23^. Although the common mechanisms of heart failure and aging are an active research topic, current research focus is still on investigating the potential anti-aging mechanisms of heart failure therapeutic drugs, and the aging-related mechanisms and potential therapeutic drugs for heart failure remain unclear^24^. Therefore, the purpose of this study was to identify and verify aging-related genes in heart failure, and to explore the mechanism and potential therapeutic drugs of aging in heart failure, in order to reveal new pathological mechanisms and cardioprotective pathways.

In this study, we first obtained 252 aging-related genes in heart failure through WGCNA and CellAge databases, and explored the biological functions and signaling pathways involved in aging genes through GO and KEGG enrichment analysis. The results show that biological processes are mainly related to histone modifications and cell cycle. Histone modifications are chemical modifications of histone amino acid residues, which can regulate gene expression without changing the DNA sequence, including methylation, acetylation, ubiquitination, etc^25^. Histone modifications are dynamically regulated under cardiac stress, leading to heart failure through compensatory or maladaptive transcriptome reprogramming^26^. Studies have shown that histone acetylation regulators can affect processes such as cardiomyocyte hypertrophy, apoptosis, fibrosis, oxidative stress, and inflammation, and exert cardioprotective effects^27^. Regulation of histone methylation and acetylation modifiers serves as a bridge between signaling and downstream gene reprogramming, and regulation of their levels helps define the epigenetic landscape required for correct cardiomyocyte function^28^. However, in aging individuals, due to histone loss, abnormal modifications, and accumulation of mutations, the strict regulation of histone modifications begins to disintegrate, disrupting tissue homeostasis and regeneration^29^. Given the reversibility of epigenetic regulation, epigenetic modifiers hold exciting promise in both delaying aging and treating heart failure^30^. Loss of cardiac contractile substrate and limited myocyte regenerative capacity are major contributors to poor outcomes in heart failure^31^. The heart is an organ with poor regenerative capacity, and it is difficult for cardiomyocytes to re-enter the cell cycle for regeneration and repair. Studies have shown that a combination of cell cycle regulators can induce stable cytokinesis in adult postmitotic cells and significantly improve cardiac function after acute or subacute myocardial infarction^32^. Additionally, forcing cardiomyocytes to proliferate while minimizing the oncogenic potential of cell cycle factors using novel transient and cardiomyocyte-specific viral constructs may reduce arrhythmias or systemic tumorigenesis while sustainably improving cardiac function^33^. Aging requires cell cycle arrest in response to damaging stimuli, and therefore, cell cycle modulators may have better efficacy in treating heart failure in the aging population^34^.

KEGG enrichment results show that the aging genes in heart failure mainly involve cellular senescence, proteoglycans in cancer, cell cycle, microRNAs in cancer, c-type lectin receptor signaling pathway, PI3K-Akt signaling pathway, and signal pathways related to cancer diseases. Cellular aging, characterized by an irreversible arrest in the cell cycle induced by stress, markedly impairs various cellular functions, including homing, proliferation, migration, and differentiation^35^. Beyond the hallmarks of DNA damage, endoplasmic reticulum stress, and mitochondrial dysfunction, senescent cardiomyocytes also exhibit an age-related secretory phenotype. This phenotype involves the release of pro-inflammatory cytokines, chemokines, and matrix-degrading enzymes, which detrimentally influence the myocardial microenvironment and neighboring healthy cardiomyocytes, exacerbating cardiac remodeling and failure^36^. Therefore, mitigating the decline in cardiac function in aging organisms necessitates not only the activation of maintenance and repair mechanisms but also prioritizing the induction of apoptosis in senescent cells, a strategy that holds promise as a therapeutic approach^37,38^. Senescent cells frequently exhibit activation of the PI3K-Akt signaling pathway, a phenomenon not observed in younger cells^39^. Interestingly, reducing AKT and ERK activation has proven effective in extending lifespan in Drosophila^40^. However, this poses a paradox, as the amelioration of myocardial fibrosis and protection of cardiac cells often entail activating the PI3K-Akt pathway^41–43^. Thus, the challenge lies in striking a balance between mitigating heart failure and aging when modulating the PI3K-Akt signaling pathway, a key area for future research.Heart failure-associated aging genes have been found to be significantly enriched in pathways commonly implicated in cancer. This correlation may stem from the intricate relationship between cellular aging, the cell cycle, and cancer. While aging naturally serves as a deterrent against tumorigenesis, senescent cells, both malignant and non-malignant, under certain conditions, can paradoxically adopt tumor-promoting characteristics^44,45^. Consequently, therapies that promote aging processes present as a viable strategy in cancer treatment^46^. Nonetheless, the multifaceted role of aging in diverse physiological and pathological contexts necessitates a careful consideration of the cardiac implications of pro-aging therapies in cancer patients. Furthermore, the role of cell division in cancer progression is critical; inaccuracies during this process can lead to chromosomal content variations and aneuploidy, thereby contributing to oncogenesis^47^. Research indicates a reduction in cell proliferation within the aging transcriptome, contrasted by a shift towards heightened cell division in the cancer transcriptome^48^. This observation suggests that a strategic, sequential application of pro-aging therapy followed by anti-aging treatment may offer a balanced approach, mitigating organ-specific burdens in cancer patients^49^.

In order to explore the key aging genes in heart failure, we used three machine learning algorithms to obtain 10 key up-regulated genes and 4 key down-regulated genes. Subsequently, we fit the 14 key genes on 15 machine learning algorithm models and 207 combinations and validated them in two independent external data sets. The results showed that the best average accuracy was 0.911, which shows that these 14 key genes can be used as aging signature genes for heart failure. This discovery paves the way for further exploration of crucial aging-related mechanisms in heart failure and the development of targeted therapeutics. Notably, the reproducibility of our findings was corroborated by their consistency across two separate and independent external datasets.

Our study identified 10 key up-regulated genes predominantly involved in cell cycle regulation, programmed cell death, and immune response. Among these, CDKN1B and SPN1 emerge as vital regulators of cell cycle progression. CDKN1B acts as a principal driver of cell division and plays a crucial role in restraining abnormal cell proliferation^50^. SPN1, associated with meiotic spindles, has been observed to induce metaphase arrest and chromosomal instability upon overexpression^51^.In the realm of programmed cell death, genes like ITPK1, MAVS, RAF1, and XAF1 play diverse roles. ITPK1 intervenes in TNF-α-induced apoptosis by disrupting the activation of the TNFRSF1A-associated death domain and is implicated in the oligomerization and localization of activated pMLKL to the cell membrane, thereby modulating necroptosis^52,53^. MAVS, while offering apoptosis resistance, also mediates the recruitment of NLRP3 to mitochondria, triggering the activation of the NLRP3 inflammasome and consequent pyroptosis^54,55^. RAF1 acts as a critical link within the MAPK/ERK cascade, determining cell fate across a spectrum of processes such as growth, proliferation, migration, differentiation, and survival^56^. It also safeguards cells from apoptosis through NF-kappa B activation and its mitochondrial translocation to bind with BCL2^57^. XAF1, in synergy with TNF-α, induces apoptosis and is involved in trophoblast cell apoptosis^58^.Furthermore, TLR3 plays a pivotal role in both innate and adaptive immunity. It operates via the TRIF/TICAM1 adapter, leading to NF-kappa B activation, IRF3 nuclear translocation, cytokine secretion, and inflammatory responses^59^.

The 4 key genes identified as down-regulated in our study play diverse roles in various biological processes. BCL6 functions as a transcriptional repressor, primarily in germinal center B cells, where it inhibits genes associated with differentiation, inflammation, apoptosis, and cell cycle regulation^60^. MEIS2, known for promoting the proliferation of cardiac myoblasts, exhibits decreased expression in aging individuals, potentially exacerbating the decline in cardiac function^61^. SMARCA2 is implicated in transcriptional activation and selective gene repression via chromatin remodeling. Research indicates that the SWI/SNF ATP-dependent chromatin remodeling complex is vital for maintaining metabolic homeostasis in adult cardiomyocytes^62^. Lastly, EIF4EBP1, which is phosphorylated in response to signals such as insulin, plays a role in the regulation of mRNA translation upon dissociation from eIF4E. This gene is also implicated in processes like autophagy and acts as a crucial effector in the mTOR signaling pathway^63,64^.

After obtaining the key aging genes of heart failure, we tried to search for potential drug molecules that can combat the development of heart failure in aging patients through the DSigDB database. The results show that Arsenenous acid, cyclophosphamide, lovastatin, Rimonabant hydrochloride, Sorafenib, and alvespimycin can interfere with some key aging genes. These drugs are mainly divided into anti-tumor drugs and lipid-lowering drugs. Arsenenous acid, cyclophosphamide, Sorafenib, and alvespimycin are predicted to be anti-tumor drugs, which may be related to key genes involved in cell cycle and programmed cell death. However, these anti-tumor drugs are generally cardiotoxic and pro-aging, which is a shortcoming of the DSigDB database^65,66^.Rimonabant, a cannabinoid receptor-1 (CB1) antagonist, shows promise in cardiovascular disease prevention^67^. Research indicates that Rimonabant not only mitigates doxorubicin-induced cardiotoxicity but also effectively reduces inflammation and oxidative stress in the aging heart^68,69^. Furthermore, it combats aging-related insulin resistance and metabolic dysfunction, reverses obesity phenotypes in aged mice, and partially restores skeletal muscle function^70,71^. These findings suggest Rimonabant’s potential in delaying aging, enhancing metabolic health, and safeguarding cardiac function.Lovastatin, known as an HMG-CoA reductase inhibitor, is widely used clinically for cholesterol reduction and vascular atherosclerosis management. However, emerging studies reveal that beyond its cholesterol-lowering capabilities, lovastatin possesses anti-aging and anti-cancer properties^72,73^. Additionally, the dedifferentiating effects of statins may alleviate myocardial fibrosis in patients predisposed to heart failure^74^. Consequently, the multifaceted mechanisms and therapeutic applications of statins like lovastatin in the realms of heart failure and aging warrant further exploration.

The novelty of our research is as follows. First, we identified common genes in heart failure and aging through WGCNA and CellAge databases. Secondly, we identified key aging genes in heart failure through 3 machine learning algorithms. Notably, we fit 14 key genes on 15 machine learning algorithm models and 207 combinations and validated them in two independent external data sets. The results show that these 14 key genes can be used as aging signature genes in heart failure, which will help to further search for key aging-related mechanisms in the process of heart failure and develop specific drugs. Finally, rimonabant and lovastatin, which we found through the DSigDB database, have the potential to delay aging and protect the heart.

Despite the contributions of this study, certain limitations must be acknowledged. Primarily, the correlation between the observed increases in mRNA levels and corresponding changes in protein expression remains uncertain. This is particularly relevant as the execution of numerous biological functions hinges on post-translational modifications. Furthermore, there is a possibility that the CellAge and DSigDB databases might have overlooked some critical genes during their screening processes. In future research endeavors, should resources allow, we plan to incorporate experimental designs that assess protein levels and drug efficacy to substantiate and refine our conclusions more robustly.

## CONCLUSION

We performed bioinformatics analysis on the GEO dataset to explore the underlying molecular mechanisms and key genes of heart failure and aging. Through three machine learning algorithms: LASSO, RF and SVM-RFE, we identified 14 key aging genes in heart failure. After fitting 15 machine learning algorithm models and 207 combinations, and validating them in two independent external data sets, we determined that these 14 key genes can serve as aging signature genes for heart failure. Our exploration via the DSigDB database revealed rimonabant and lovastatin as promising agents capable of decelerating aging processes and offering cardiac protection. Collectively, these insights pave the way for enhanced understanding of aging-related mechanisms in heart failure and could inform the development of targeted therapeutic interventions.

## Acknowledgements

Not applicable.

## Authors’ contributions

YDY conducted statistical analysis and drafted the article. LW and WJH contributed to picture processing and article reviewing. YTX reviewed and proofread the article. XJL and YL provided effective scientific suggestions and supervision and created the final revision of the manuscript. All authors read and approved the final manuscript.

## Consent for publication

Not applicable.

## Ethics approval and consent to participate

Not applicable.

## Availability of data and materials

Publicly available datasets were analyzed in this study. This data can be found here: GSE57338; GSE5406; GSE16499.

